# *Drosophila* Trpm mediates calcium influx during egg activation

**DOI:** 10.1101/663682

**Authors:** Qinan Hu, Mariana F. Wolfner

**Affiliations:** Department of Molecular Biology and Genetics, Cornell University, Ithaca, NY 14853

**Keywords:** egg activation, oogenesis, embryogenesis, Trpm, germline specific CRISPR/Cas9, *Drosophila melanogaster*

## Abstract

Egg activation is the process in which mature oocytes are released from developmental arrest and gain competency for embryonic development. In *Drosophila* and other arthropods, eggs are activated by mechanical pressure in the female reproductive tract, whereas in most other species, eggs are activated by fertilization. Despite the difference in the trigger, *Drosophila* shares many conserved features with higher vertebrates in egg activation, including a rise of intracellular calcium in response to the trigger. In *Drosophila*, this calcium rise is initiated by entry of extracellular calcium due to opening of mechanosensitive ion channels and initiates a wave that passes across the egg prior to initiation of downstream activation events. Here, we combined inhibitor tests, germline specific RNAi knockdown, and germline specific CRISPR/Cas9 knockout to identify the Transient receptor potential (TRP) channel subfamily M (Trpm) as a critical channel that mediates the calcium influx and initiates the calcium wave during *Drosophila* egg activation. We observed reduced egg hatchability in *trpm* germline knockout mutant females, although eggs were able to complete some egg activation events including cell cycle resumption. Since the mouse Trpm ortholog was recently reported also to be involved in calcium influx during egg activation and in further embryonic development our results suggest that calcium uptake from the environment via TRPM channels is a deeply conserved aspect of egg activation.

**Significance:** A rise in intracellular free calcium is a conserved feature of the egg-to-embryo transition in almost all animals. In *Drosophila*, as in vertebrates, the rise starts at one end of the egg, and then travels across the egg in a wave. The *Drosophila* calcium rise is mediated by an influx of calcium, due to the action of mechanically-gated ion channels. Here we identify the ion channel that is critical for the calcium entry as TRPM. TRPM is the ortholog of the channel recently shown to mediate the post-fertilization calcium influxes needed to sustain calcium oscillations in fertilized mouse eggs, suggesting a deep homology – despite species differences in the trigger for egg activation.

## Introduction

In almost all animals, mature oocytes are arrested in meiosis at the end of oogenesis and require an external trigger to be activated and transition to start embryogenesis. This “egg activation” involves multiple events that transition eggs to embryogenesis, including meiosis resumption and completion, maternal protein modification and/or degradation, maternal mRNA degradation or translation, and egg envelope changes (reviewed in (1–5)).

Triggers of egg activation vary across species. In vertebrate and some invertebrate species, fertilization triggers egg activation. However, changes in pH, ionic environment, or mechanical pressure can also trigger egg activation in other invertebrate species (reviewed in (1)). A conserved response to these triggers is a rise of intracellular free Ca^2+^ levels in the oocyte. This calcium rise is due to influx of external calcium and/or release from internal storage, depending on the organism (reviewed in (5)). The elevated Ca^2+^ concentration is thought to activate Ca^2+^-dependent kinases and/or phosphatases, which in turn change the phospho-proteome of the activated egg, initiating egg activation events (reviewed in (3)).

*Drosophila* eggs activate independent of fertilization and the trigger is mechanical pressure. When mature oocytes exit the ovary and enter the lateral oviduct, they experience mechanical pressure from reproductive tract tract. As the oocytes swell due to the influx of oviductal fluid, their envelopes become taut (6). *Drosophila* oocytes can be activated *in vitro* by incubation in a hypotonic buffer, although some egg activation events do not proceed completely normally *in vitro* (7, 8). Intracellular calcium levels rise in oocytes occur egg activation, as observed with the calcium sensor GCaMP. This calcium rise takes the form of a wave that starts at the oocyte pole(s) and traverses the entire oocyte (9, 10). Initiation of this calcium wave requires influx of external Ca^2+^, as chelating external Ca^2+^ in *in vitro* egg activation assays blocks the calcium wave and egg activation. Propagation of the calcium wave relies on the release of internal Ca^2+^ stores, likely through an Inositol 1,4,5-trisphosphate (IP_3_) mediated pathway, as knocking down the endoplasmic reticulum (ER) calcium channel IP3 receptor (IP_3_R) prevents propagation of the calcium wave (9).

How mechanical forces trigger calcium entry during *Drosophila* egg activation was unknown. However, the lack of initiation of a calcium wave in the presence of Gd^3+^, an inhibitor of mechanosensitive ion channels (11), and N-(p-Amylcinnamoyl) anthranilic acid (ACA), an inhibitor of TRP-family ion channels (12), suggested that TRP family ion channels (reviewed in (13)) are likely involved (9). Further supporting this idea is the recent discovery that a TRP family channel, TRPM7, is needed for calcium influx that leads to calcium oscillations in activating mouse eggs (14). The *Drosophila* genome encodes 13 TRP family channels (reviewed in (15)), but according to RNAseq data, only 3 (Painless, Trpm, and Trpml) are expressed in the ovary (reviewed in (2)). We used specific inhibitors, existing mutants, germline-specific RNAi knockdown, and new knockouts that we created with CRISPR/Cas9 to screen these 3 candidates for their roles in the initiation of the calcium wave. We found that Trpm, the single *Drosophila* ortholog of mouse TRPM7, mediates the calcium wave initiation, whereas the other two TRP channels are not necessary to initiate the calcium wave.

## Results

### Painless and Trpml are not essential for the calcium wave initiation or propagation

The three TRP channels (Painless (Pain), Trpm and Trpml) expressed in the *Drosophila* ovary and are candidates for involvement in calcium wave initiation. To examine the roles of these channels in calcium wave during egg activation, we performed *in vitro* activation assays on mature oocytes dissected from wildtype, mutant, and/or knockdown females, and imaged the calcium waves with fluorescent microscopy (see Materials and Methods).

Among the three candidates, Pain is reported to display mechanosensitivity (16). To examine the role of Pain, we made a CRISPR knockout line, *pain^TMΔ^*, by deleting the region that encodes Pain’s transmembrane domains. Since *pain* loss-of-function mutants are viable (17), we were able to examine the calcium wave phenotype in oocytes from homozygous *pain^TMΔ^* females that also carried *a nos-GCaMP6m-attP2* transgene. The frequency of observing a calcium wave *in vitro* in mature oocytes of homozygous *pain^TMΔ^* (n=33/48) females did not differ significantly from that of heterozygous controls (**Fig.1A, Movie S1-2**). The propagation rate of the calcium wave, as measured at its wave-front, was also not significantly different (**Fig.1B**).

**Fig. 1.**
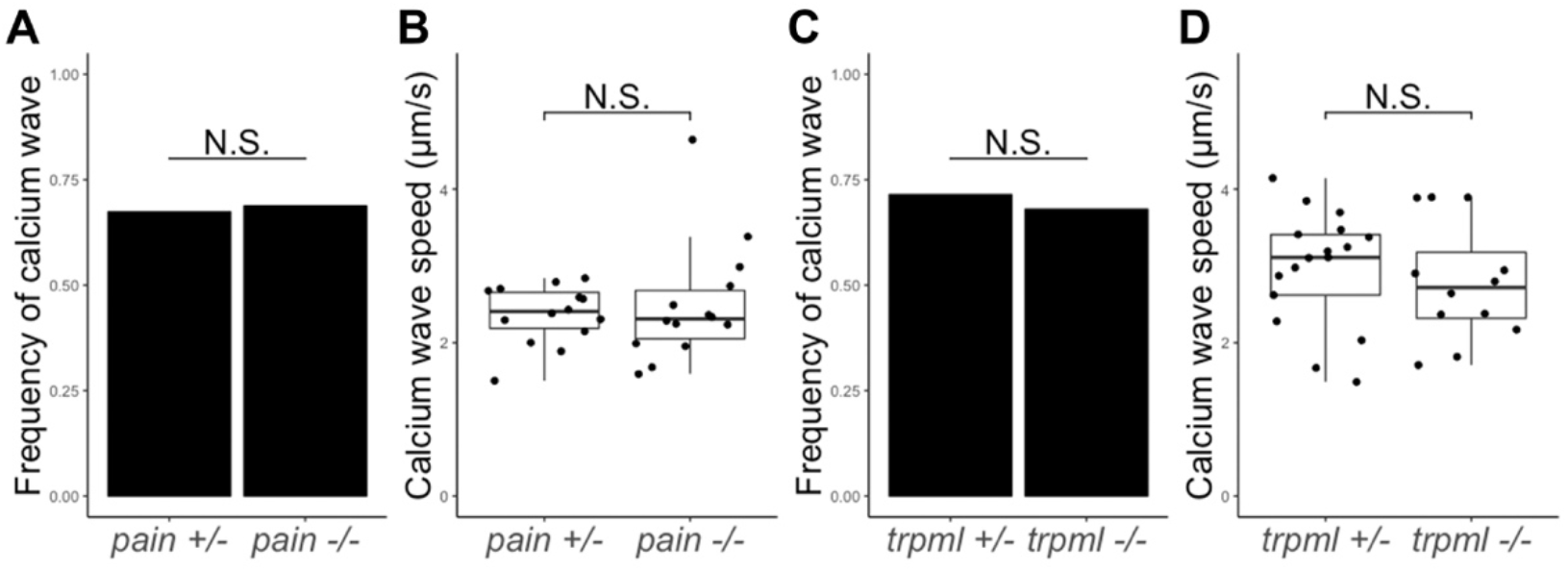
*pain* and *trpml* are not essential for calcium wave initiation or propagation. **(A)** The frequency of observing a calcium wave in *in vitro* activation assays does not differ between control *(pain^TMΔ^/+*, n=33/49) and *pain* null mutant *(pain^TMΔ^/pain^TMΔ^*, n=33/48), p=1; **(B)** The propagation speed of the calcium wave does not differ between control (2.37±0.38 μm/s) and *pain* null mutant (2.49±0.78 μm/s), p=0.59; **(C)** The frequency of observing a calcium wave in *in vitro* activation assays does not differ between control *(trpml*^1^/+, n=17/21) and *trpml* null mutant *(trpml^1^/trpml^1^*, n=15/25), p=0.22; **(D)** The propagation speed of the calcium wave does not differ between control (2.97±0.74 μm/s) and *trpml* null mutant (2.78±0.77 μm/s), p=0.51.

To examine the role of Trpml, we tested an existing null mutant, *trpml^1^* (18). We crossed into *trpml^1^* the *nos-GCaMP6m-attP40* transgene and examined the calcium wave phenotype in mature oocytes from homozygous mutants or heterozygous controls. Again, we did not find a difference in the calcium wave initiation frequency (**Fig.1C, Movie S3-4**) or in the calcium wave propagation rate between oocytes from homozygous mutant and those from heterozygous control females (**Fig. 1D**).

Thus, neither Pain or Trpml is essential for initiation or propagation of the calcium wave. These channels thus are either not involved in calcium wave initiation, or they function redundantly with other channels.

### Inhibitor and RNAi perturbations of Trpm inhibit calcium wave initiation

To test whether Trpm is needed for the calcium wave, we first examined the effect of known inhibitors of this channel. 10 mM Mg^2+^ is reported to inhibit several TRP channels including mouse TRPV3 (19), TRPM6, TRPM7 (20) and *Drosophila* Trpm (21). In the presence of 10 mM Mg^2+^ in IB and RB, the frequency of observing a calcium wave *in vitro* was significantly reduced (**Fig.2A, B&E, Movie S5**). We also tested the effect of NS8593, a specific inhibitor of mouse TRPM7 channels (22). 100 μM NS8593 in IB and RB completely blocked calcium wave initiation *in vitro* (**Fig.2C&F, Movie S6**).

**Fig. 2.**
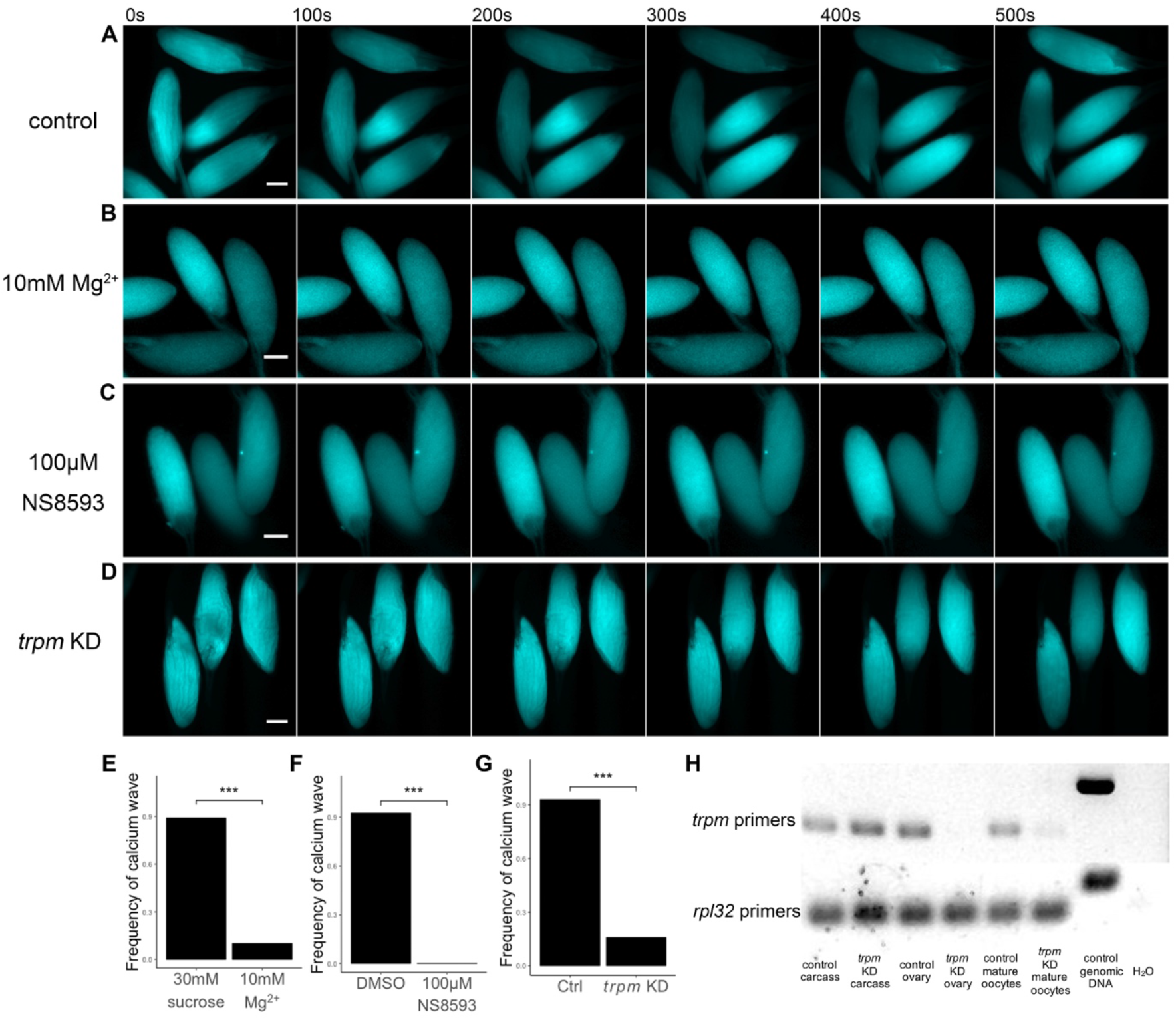
Disrupting *trpm* function reduces calcium wave frequency *in vitro*. **(A-C)** Representative calcium waves, or lack thereof, in *matα4-GAL-VP16; UASp-GCaMP3* stage 14 oocytes in *in vitro* activation assay. Oocytes were incubated in: **(A)** unmodified IB and RB; **(B)** IB and RB with 10mM of MgCl_2_; **(C)** IB and RB with 100 μM NS8593; **(D)** Representative pictures showing lack of calcium waves in stage 14 oocytes from *trpm* germline knockdown females *(matα4-GAL-VP16/UAS-trpm^RNAli^; UASp-GCaMP3)* incubated in unmodified IB and RB. All scale bars=50 μm; **(E-G)** Quantification of calcium wave frequency in oocytes as in **(B-D)**: **(E)** n=16/18 for control (IB and RB with 30mM sucrose to provide the same osmolarity change as 10mM MgCl_2_), n=2/20 for 10mM MgCl_2_, p=5.69×10^−6^; **(F)** n=27/29 for control (DMSO), n=0/29 for 100 μM NS8593, p=2.15×10^−11^; **(G)** n=26/28 for control *(matα4-GAL-VP16; UASp-GCaMP3)*, n=11/70 for *trpm* germline knockdown mutants *(matα4-GAL-VP16/UAS-trpm^RNAi^; UASp-GCaMP3*), p=5.74×10^−12^; **(H)** RT-PCR quantification of *trpm* germline knockdown. Normalized expression level of *trpm* was calculated by normalizing *trpm* band intensity with that of ribosome protein gene *rpl32*. Transcription level of *trpm* is decreased by >90% in knockdown ovaries and mature oocytes. Both sets of primers flank an exon-exon junction and will amplify a bigger band with genomic DNA as templates. This ensures that the bands we quantified were amplified from cDNA only.

Since *trpm* homozygous mutants are lethal before adult stage (23), we next tested the role of *trpm* in calcium wave initiation using germline-specific RNAi. We crossed *matα4-GAL-VP16; UASp-GCaMP3* to *UAS-trpm^RNAi^*. The female offspring expressed both GCaMP3 and *trpm* shRNA in the germline. We examined the calcium waves in *in vitro* activation assays with mature oocytes from these females. Oocytes form *trpm* germline knockdown females displayed a significantly lower frequency of calcium waves *in vitro* (**Fig.2D&G, Movie S7**). To validate the efficiency of germline knockdown, we performed RT-PCR with ovary mRNA from these females. RT-PCR results showed that more than 90% of *trpm* transcripts were removed in germline knockdown females compared to control (**Fig.2H**).

Inhibitors of Trpm and germline knockdown of *trpm* both reduced the frequency of observing calcium wave in *in vitro* egg activation assays. Thus, our data strongly suggested that Trpm is necessary for calcium influx during *Drosophila* egg activation.

### Germline specific CRISPR/Cas9 knockout of *trpm* inhibits calcium wave initiation

To further validate the role of Trpm in the initiation of the calcium wave, we attempted to perform germline specific biallelic knockout of *trpm* using the CRISPR/Cas9 system. We designed three sets of gRNAs targeting *trpm* (see Materials and Methods). We used the “Cas9-LEThAL” (24) method to evaluate the efficiency of our gRNA expression constructs. Among them, gRNA set 3 displayed the highest efficiency (**Table.S1**). Using this set of gRNAs, we made a dual-gRNA transgenic line based on a polycistronic gRNA design (*U6:3-tRNA-gRNA1-tRNA-gRNA2, gRNA-trpm^1^*) and a dual transcription unit design *(CR7T-gRNA1-U6:3-gRNA2, gRNA-trpm^2^*), both ubiquitously expressing the gRNAs.

We crossed these two transgenic fly strains to *nos-Cas9; matα4-GAL-VP16; UASp-GCaMP3* to achieve germline specific knockout of *trpm* and calcium visualization at the same time. By sequencing amplicons generated by single oocyte RT-PCR, we confirmed the presence of CRISPR/Cas9 generated indels (**Fig.S2**, see Supplemental Materials and Methods). With *gRNA-trpm^1^*, 34% of the female offspring displayed defects in ovary morphology (**Fig.S3A-B**). With *gRNA-trpm^2^*, 96% of the female offspring displayed similar defects (**Fig.S3C**). These data showed that *trpm* is required for early female germline development. Since Trpm is involved in maintaining cation homeostasis in cellular environments and tissue development (21, 23), we hypothesize that early knockout of *trpm* by *nos-Cas9* interferes with normal germline development. The less-than-100% frequency of this phenotype is likely due to lack of 100% efficiency in generating biallelic null mutations of *trpm* in the germline with less efficient gRNA expression constructs.

Some *nos-Cas9; gRNA-trpm^1^* females had ovaries with grossly normal morphology, perhaps due to null mutations generated at both *trpm* alleles after a critical development point. We tested their mature oocytes for activation *in vitro*. We observed that the frequency of calcium wave initiation was significantly reduced in oocytes from these females (**Fig.3A, B&D, Movie S8**). Since efficient CRISPR/Cas9 mediated *trpm* knockout caused ovary development defects, flies with normal-looking ovaries may have less efficient *trpm* biallelic knockout in the germline, which could explain the incomplete elimination of calcium waves in oocytes from these flies.

**Fig. 3.**
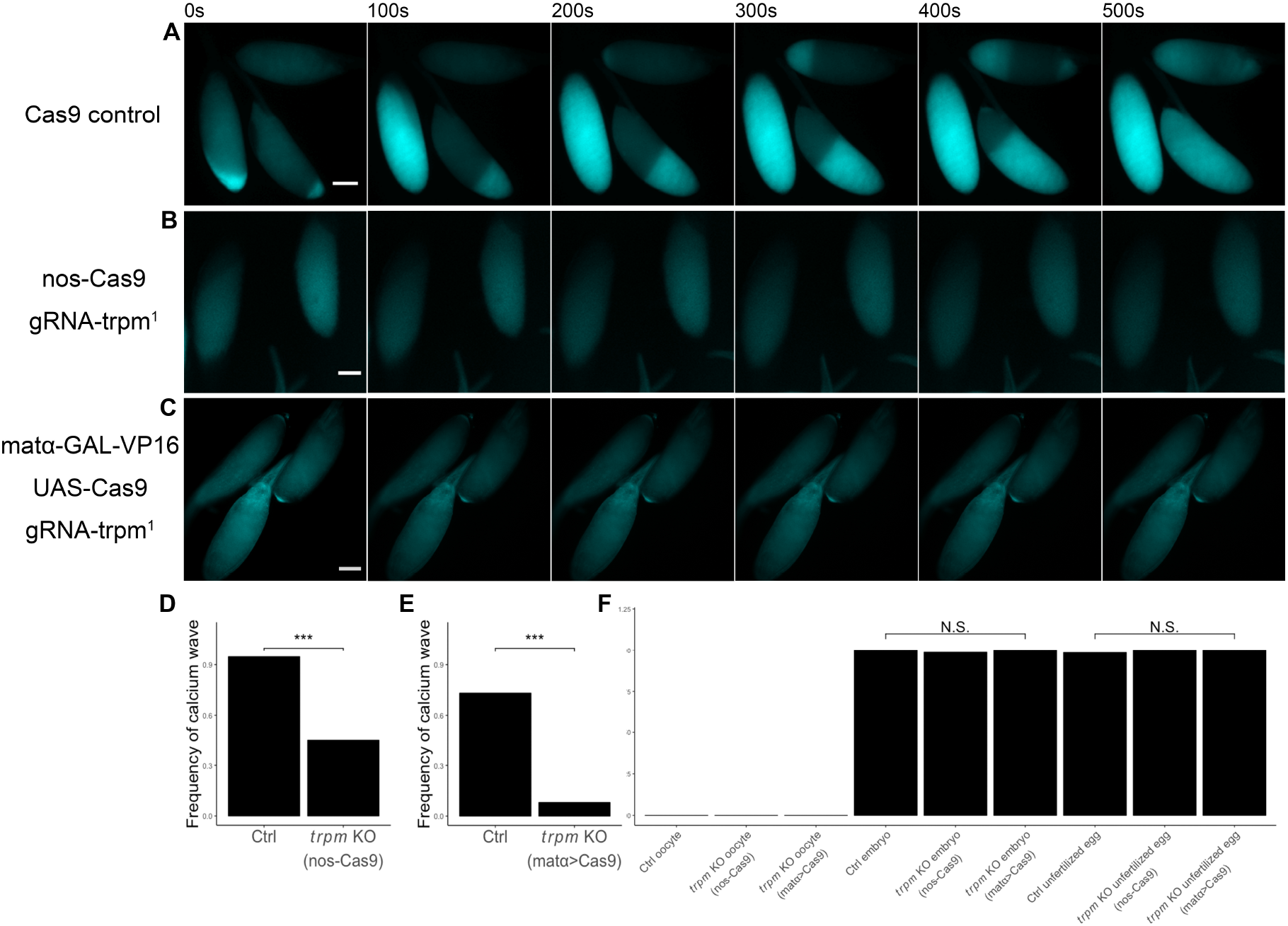
*trpm* germline specific CRISPR knockout reduces calcium wave frequency and egg hatchability. **(A-C)** Representative calcium waves, or lack thereof, in *in vitro* activation assay. Stage 14 oocytes were dissected from: **(A)** *nos-Cas9; matα4-GAL-VP16; UASp-GCaMP3* control; **(B)** *nos-Cas9; matα4-GAL-VP16; UASp-GCaMP3/gRNA-trpm^1^;* **(C)** *matα4-GAL*- VP16/UAS-GCaMP6s; *gRNA-trpm^1^/UAS-Cas9*. All scale bars=50 μm; **(D-E)** Quantification of calcium wave frequency in oocytes as in **(B-C): (D)** n=36/39 for control *(nos-Cas9; matα4-GAL-VP16; UASp-GCaMP3)*, n=38/87 for *trpm* germline knockout *(nos-Cas9; matα4-GAL-VP16; UASp-GCaMP3/gRNA-trpm^1^)*, p=4.64×10^−7^; **(E)** n=20/27 for control *(matα4-GAL-VP16/UAS-GCaMP6s; UAS-Cas9)*, n=5/62 for *trpm* germline knockout (*matα4-GAL-VP16/UAS-GCaMP6s*; *gRNA-trpm^1^/UAS-Cas9*), p=3.68×10^−11^; **(F)** Resistance to 50% bleach by mature oocytes, 1-5h embryos and unfertilized eggs from control *(nos-Cas9-attP2)* and *trpm* germline CRISPR knockout *(nos-Cas9-attP2/gRNA-trpm^1^* and *matα4-GAL-VP16; UAS-Cas9/ gRNA-trpm^1^)* females. All pre-activation oocytes are vulnerable to bleach. 1-5 h embryos and unfertilized eggs from both control and *trpm* germline knockout mutants displayed resistance to bleach treatment. Control oocyte n=132, *trpm* KO *(nos-Cas9)* oocyte n=152, *trpm* KO *(matα>Cas9)* oocyte n=119, control embryo n=76, *trpm* KO *(nos-Cas9)* embryo n=73, *trpm* KO *(matα>Cas9)* embryo n=64, control unfertilized egg n=66, *trpm* KO *(nos-Cas9)* unfertilized egg n=58, *trpm* KO *(matα>Cas9)* unfertilized egg n=71.

To bypass the critical stage in early oogenesis that may require *trpm* function, we knocked out *trpm* at only later stages of oogenesis by crossing the *UAS-GCaMP6s; gRNA-trpm^1^* strain to the *matα4-GAL-VP16; UAS-Cas9* strain. Since *matα4-GAL-VP16* does not drive expression of UAS constructs until mid to late oogenesis (25), female offspring of this cross will initiate germline knockout of *trpm* at later stages of oogenesis. None of these germline knockout females displayed gross ovary morphology defects (**Fig.S3D-E**), suggesting that *trpm* knockout occurred after a critical stage in early oogenesis. We examined the calcium wave phenotype of mature oocytes from these females in *in vitro* egg activation assays. We observed a significant decrease in the frequency of calcium wave initiation (**Fig.3C&E, Movie S9**).

Taken together, these results further supported that Trpm is required for calcium wave initiation. In addition, our results also suggested that Trpm plays essential roles in early oogenesis.

### Egg activation events occur in *trpm* knockout eggs *in vivo*

We next asked whether egg activation requires normal *trpm* function. Almost all 1-5 h embryos from *trpm* germline knockout females displayed resistance to 50% bleach, an indicator of vitelline membrane crosslinking after egg activation (26) (**Fig.3F**). This suggests that egg envelope hardening, one aspect of egg activation, still occurs in embryos from *trpm* germline knockout females.

Since bleach resistance does not always completely reflect the state of egg activation (8), we examined whether embryos from *trpm* germline knockout females were able to enter mitosis, implying successful completion of meiosis and fertilization. Surprisingly, anti-Tubulin and DAPI staining of 1-5 h old embryos laid by *trpm* germline knockout females mated with ORP2 wildtype males showed they had all undergone early embryo mitosis (n=88 for *nos-Cas9* knockout mutants, n=36 for *matα4-GAL-VP16 > UAS-Cas9* knockout mutants, **Fig.4A**). This indicated that cell cycles can resume in embryos from *trpm* germline knockout females mated with wildtype males.

**Fig. 4.**
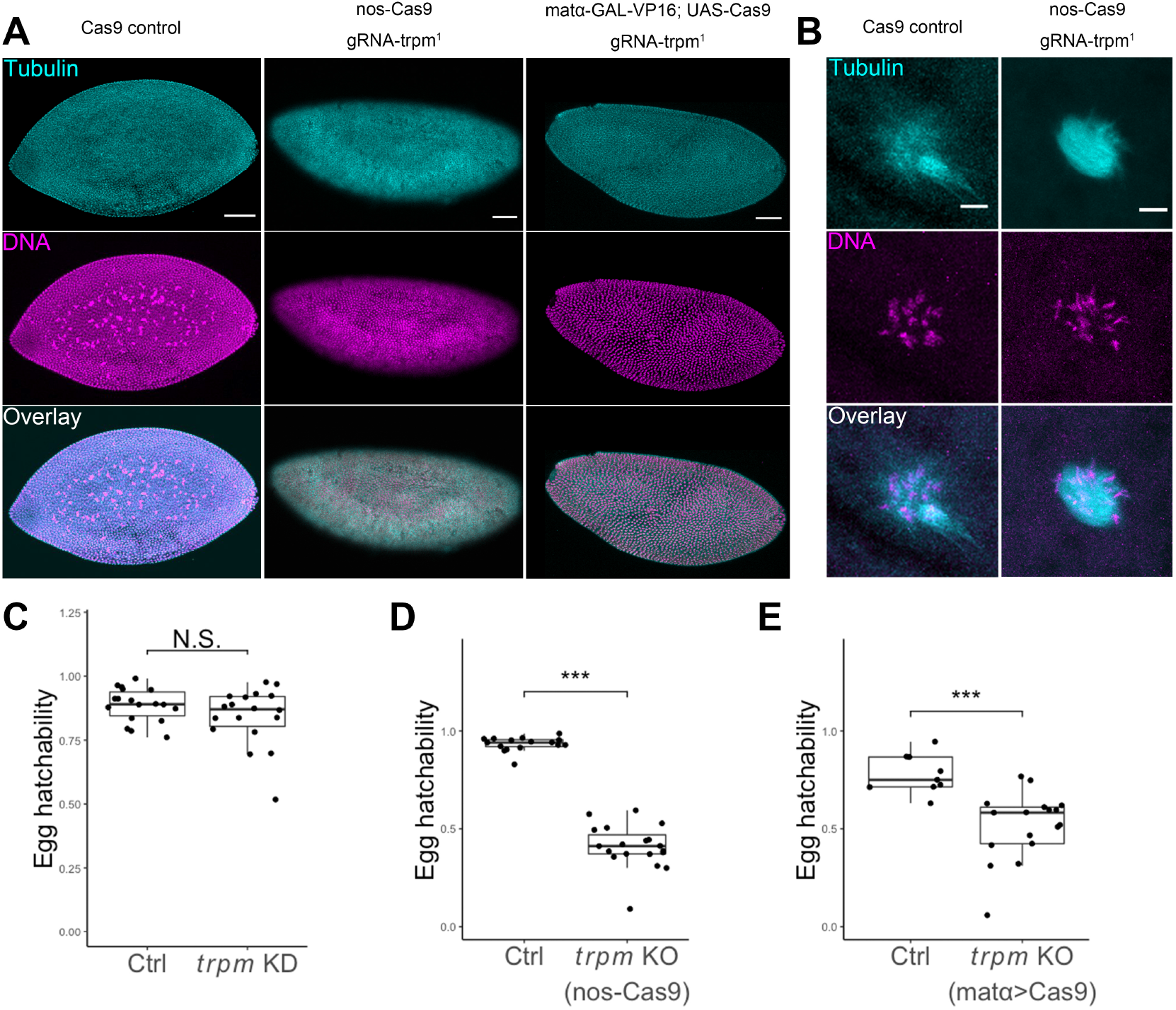
Cell cycle resumption and vitelline membrane crosslinking occur normally in eggs from *trpm* germline knockout females. **(A)** Representative pictures of anti-Tubulin and DAPI staining of 1-5 h embryo laid by control and *trpm* germline CRISPR knockout females. Focal planes vary across samples. All embryos from *trpm* germline knockout females have started mitosis and progressed to multicellular stages (n=88 for knockout with *nos-Cas9*, n=36 for knockout with *matα4-GAL-VP16 > UAS-Cas9)*. Scale bars=50 μm. **(B)** Representative pictures of anti-Tubulin and DAPI staining of unfertilized eggs laid by control *(nos-Cas9-attP2*, n=14) and *trpm* germline knockout *(nos-Cas9-attP2/ gRNA-trpm^1^*, n=20) females. All eggs from both control and germline knockout females have completed meiosis, indicated by production of normal polar body. Scale bars=5 μm. **(C)** *trpm* germline RNAi *(matα4-GAL-VP16/UAS-trpm^RNAi^; UASp-GCaMP3*, n=18) does not significantly affect female egg hatchability relative to control *(matα4-GAL-VP16; UASp-GCaMP3*, n=18), p=0.15; **(D)** *trpm* germline CRISPR knockout *(nos-Cas9-attP2/gRNA-trpm^1^*, n=21) significantly reduces female egg hatchability relative to control *(nos-Cas9-attP2*, n=16), p=2.08×10^−15^; **(E)** *trpm* germline CRISPR knockout *(matα4-GAL-VP16; UAS-Cas9/gRNA-trpm^1^*, n=17) significantly reduces female egg hatchability relative to control *(matα4-GAL-VP16 > UAS-Cas9*, n=9), p=5.13×10^−5^.

We then asked if egg hatchability is affected by *trpm* germline perturbations. Germline knockdown of *trpm* did not impair egg hatchability (**Fig.4C**). Since this knockdown, despite being highly efficient, did not completely abolish *trpm* function in oocytes, we examined the hatchability of eggs from *trpm* germline knockout females. Germline *trpm* knockout females displayed significantly reduced egg hatchability compared to control, with either *nos-Cas9* (**Fig.4D**) or *matα4-GAL-VP16 > UAS-Cas9* (**Fig.4E**) to achieve germline knockout, suggesting that the lack of maternal Trpm function or a calcium wave compromised development events after initiation of the embryos’ mitotic phase. Since the fathers of these embryos were wildtype, a wildtype *trpm* allele was present in the embryos. Thus, defects in hatchability must be due to the lack of maternal *trpm* product, because after zygotic genome activation, these embryos will express wildtype Trpm.

Taken together, these observations indicated that fertilized eggs are able to resume cell cycles and harden egg envelopes in the absence of Trpm function. The reduced hatchability of eggs laid by *trpm* germline knockout females suggested that lack of oocyte Trpm function or a calcium wave affects embryogenesis after egg activation.

### Sperm is not an alternative trigger of egg activation in the absence of Trpm function

Because fertilized eggs from *trpm* germline knockout females proceeded to mitotic stages, we wondered whether sperm could act as an alternative trigger for calcium rise and egg activation (reviewed in (1)) in *Drosophila*. We mated control and *trpm* germline knockout females to spermless males (27). Eggs laid by females after the mating would experience normal egg-activating environment *in vivo* but remain unfertilized. We then examined if meiosis resumption and vitelline membrane crosslinking occur in unfertilized eggs from *trpm* germline knockout females. We observed 100% resistance to 50% bleach in these unfertilized eggs (**Fig.3F**). We also observed normal production of polar bodies in all unfertilized eggs from control (n=14) and *trpm* germline knockout females (n=20) (**Fig.4B**). These results indicated that vitelline membrane crosslinking and meiosis resumption still occur in eggs from *trpm* germline knockout mutants independent of sperm, ruling out the possibility that sperm could activate eggs in the absence of Trpm function.

## Discussion

Egg activation is an essential step for oocytes to transition to embryogenesis. In all species studied to date, egg activation involves a rise in intracellular calcium. In *Drosophila*, this calcium rise takes the form of a wave that is triggered by mechanical pressure. The initiation of this calcium wave requires influx of calcium from the environment through mechanosensitive TRP family channel(s) (9).

Here, we determined that Trpm channel is necessary for this calcium wave. Of the three TRP channels expressed in ovaries (Pain, Trpm and Trpml), only impairment of Trpm affects the initiation of the calcium wave. Calcium wave phenotypes are normal in oocytes from *pain* or *trpml* null mutants. However, the frequency of the calcium wave is diminished in wildtype oocytes in the presence of Trpm inhibitors and in oocytes from *trpm* germline knockdown or knockout mutants. These results consistently indicated that Trpm mediates the calcium influx that initiates the calcium wave during *Drosophila* egg activation. *trpm* germline knockout females also displayed significantly decreased egg hatchability, due to defects after cell cycle resumption. The reduced hatchability suggested that maternal *trpm* function or the calcium wave is required for further embryogenesis after egg activation.

### The *Drosophila* Trpm channel plays important reproductive roles

TRP family ion channels are non-selective and respond to a wide array of environmental stimuli. *Drosophila* Trpm has been reported to play multiple roles throughout larval development, including maintaining Mg^2+^ and Zn^2+^ homeostasis (21, 23), and sensing noxious cold in larval Class III md neurons (28). However, the role of Trpm in reproduction had not been investigated because of the pupal lethality of *trpm* null mutants (23). Here, our germline specific RNAi knockdown and CRISPR/Cas9 mediated knockout revealed three novel functions of *Drosophila* Trpm: supporting early oogenesis, mediating influx of environmental calcium to initiate the calcium wave during egg activation, and maternally supporting embryonic development after egg activation.

Our previous study suggested that calcium influx during *Drosophila* egg activation is mediated through mechanosensitive ion channels (9). Both *Drosophila* Trpm and its mouse ortholog TRPM7 are reported to be constitutively active and permeable to a wide range of divalent cations (21, 29). Mouse TRPM7 is known to respond to mechanical pressure (30, 31), but further study will be needed to determine whether *Drosophila* Trpm is similarly responsive to mechanical triggers, such as those that occur during ovulation.

### Requirement for Trpm during Drosophila embryogenesis

Germline knockout of *trpm* significantly reduced the frequency of observing calcium waves in *in vitro* egg activation assays and egg hatchability. However, this reduced egg hatchability was not due to failure of cell cycle resumption during egg activation. There are two possible explanations for the reduced egg hatchability of *trpm* germline knockout females.

First, it is possible that *trpm* plays a maternal role, independent of its role in initiating the calcium wave, such that lack of maternally deposited Trpm proteins leads to defects during embryogenesis. In mouse, TRPM7 is also required for normal early embryonic development, apart from its role in calcium oscillations. Inhibition of TRPM7 function impairs pre-implantation embryo development and slows progression to the blastocyst stage (32). *Drosophila trpm* mutant lethality had been reported to occur during the pupal stage (23). However, those homozygous mutants were offspring of heterozygous mothers, and thus did not lack maternal Trpm function. Our germline specific depletion of *trpm* reveals a maternal role for Trpm in embryogenesis.

Alternatively, or in addition, it is possible that oocytes lacking Trpm do not take up sufficient Ca^2+^ from the environment to form a calcium wave, but that at least some events of egg activation can occur despite this. In mouse, an initial calcium rise is induced by sperm-delivered PLCζ via the IP3 pathway (33). Yet although sperm from PLCζ null males fails to trigger normal calcium oscillations, some eggs fertilized by those sperm develop (34). Multiple oscillations following fertilization require influx of external calcium (35), mediated by TRPM7 and Ca_v_3.2 (14). Even though these oscillations were reported to be needed for multiple post-fertilization events (36), some TRPM7 and Ca_v_3.2 double-knockout embryos still develop, albeit not completely normally (14). Together, these data suggest that egg activation can still occur in mouse with diminished intracellular calcium rises, analogous to what we see in *Drosophila* in the absence of maternal Trpm function.

Insufficient influx of calcium in the absence of Trpm function could disrupt later (but maternally-dependent) embryogenesis. The oocyte-to-embryo transition involves multiple events. In mouse egg activation, these events take place sequentially as calcium oscillations progress, with developmental progression associated with more oscillations and more total calcium signal. Some of the events start after a certain number of oscillations but require additional oscillations to complete (36). It is possible that mechanisms critical for *Drosophila* embryo development also depend on reaching a precise level of calcium. A low-level calcium rise might be sufficient to trigger some egg activation mechanisms such as vitelline membrane crosslinking and cell cycle resumption, but high-levels of calcium may be required for further progression.

### Alternative pathways for calcium influx during egg activation

Given the importance of calcium in egg activation (8, 35), we were surprised that although *trpm* knockout eggs lacked a calcium wave *in vitro, in vivo* such eggs could progress in cell cycles and even, sometimes, hatch. As discussed above, there may be insufficient calcium influx in the absence of Trpm for full and efficient development, but some egg activation events may still occur. Alternatively, it is possible that redundant mechanisms permit a sufficient calcium-level increase without producing a detectable wave form. Despite being able to trigger a series of egg activation events including meiosis resumption and protein translation (7, 8), osmotic pressure during *in vitro* activation may have different properties from mechanical pressure exerted on mature oocytes during ovulation. The latter might allow opening of other calcium channels to initiate a normal calcium rise and complete egg activation. Two channels, TRPM7 and Ca_v_3.2, are needed for the calcium oscillations following mouse fertilization, but the *Drosophila* ortholog of mouse Ca_v_3.2, Ca-α1T, is not detectably expressed in fly ovaries (37). Other unknown channels might play this redundant role *in vivo*. In this light we note that levels of basal GCaMP fluorescence varied among oocytes incubated in IB that were inhibited from forming a wave, suggesting the possibility of a calcium increase by a redundant mechanism (**Fig.S4**).

### Conserved role of TRPM channels in egg activation

We are intrigued that *Drosophila* Trpm is essential for the calcium rise at egg activation, and that its mouse ortholog, TRPM7, was recently reported to be required (along with Ca_v_3.2) for the calcium influx needed for post-fertilization calcium oscillations that are in turn required for egg activation events (14, 36). This apparent conservation in mechanisms in egg activation involving orthologous Trpm channels in a protostome *(Drosophila)* and a deuterostome (mouse) prompts us to wonder whether Trpm-mediated calcium influx is a very ancient and basal aspect of egg activation, with other more variable aspects such as sperm-triggered calcium rises being more derived, if better known, features. It is interesting in this light that a sperm-delivered TRP channel (TRP-3) has also been reported to mediate calcium influx and a calcium rise in another protostome, *C. elegans* (38).

## Materials and Methods

### DNA constructs and transgenic flies

Calcium waves were visualized by expressing GCaMP sensors in the female germline using *matα4-GAL-VP16; UASp-GCaMP3* (9), *matα4-GAL-VP16; UAS-GCaMP6s*, or *nos-GCaMP6m*. The *nos-GCaMP6m* strain was constructed by replacing the GCaMP3 coding sequence in the previously described *nos-GCaMP3-attB* construct (9) with GCaMP6m coding sequence and integrating the construct into either the attP2 or the attP40 site.

Calcium wave frequency in each experiment was compared between perturbed and control oocytes using the same calcium sensor. Control oocytes showed lower frequency of calcium waves when expressing *GCaMP6s* or *GCaMP6m* compared to *GCaMP3*, likely due to weaker expression of our *GCaMP6s* or *GCaMP6m* constructs.

To create a null allele of *pain*, we generated *pU6-chiRNA* constructs following protocols described by FlyCRISPR website (39). We generated two constructs to express sgRNAs with the following target sequences (PAM sequences are underlined):

GTCTTGCAGCTGGTTGAGT<u>CCGG</u>, GACGCAGACTTAAGTAGTTC<u>GGG</u>. These two constructs were co-injected by Rainbow Transgenic Flies, Inc into *nos-Cas9-attP2* embryos, a strain carrying a 1091bp deletion (chr2R:24922130-24923220) in *pain* was isolated and stabilized to establish the null allele strain *pain^TMΔ^* (see Supplemental Materials and Methods). To knockout *trpm*, we generated the U6:3-tRNA-gRNA1-tRNA-gRNA2 (24) and CR7T-gRNA1-U6:3-gRNA2 constructs (unpublished from Dr. Chun Han). We designed three sets of gRNAs targeting *trpm* to express in our constructs. sgRNA target sequences in the *trpm* gene are as follows (PAM sequences are underlined):

gRNA set 1: GGAACCATCGAGTTCCAGGG<u>CGG</u>; GATGTGGACACATGGCGAGG<u>AGG</u>; CTTTTGATCACCGTGCAGGG<u>CGG</u>; CTTGGACACGGAAATCTACG<u>AGG</u> gRNA set 2: GATGAGCGAGGAGGGCACGA<u>TGG</u>; ACCCATAACCAAGTTCTGGG<u>CGG</u> gRNA set 3: GACTACAGGGATGAGCGAGGAGG; GAATACCACTCCTGCCACCGCGG These constructs were injected by Rainbow Transgenic Flies, Inc into *yw, nos-phiC31; PBac{attP-9A}* strain.

### gRNA construct efficiency test

To confirm the efficiency of constructs expressing gRNAs, we employed the “Cas9-LEThAL” method (24) to validate the efficiency of our gRNA constructs. Males carrying gRNA constructs were crossed to *act-Cas9, lig4*-mutant females. Our gRNA set 3 construct caused 100% lethality in *lig4* mutant male offspring while the other two sets did not (**Table.S1**), suggesting it has the highest efficiency.

### Fly strains and maintenance

All *Drosophila* strains and crosses were maintained or performed on standard yeast-glucose-agar media at 25C° (29C° for all crosses involving the GAL4/UAS system to enhance its efficiency (40)) on a 12/12 light/dark cycle. The following fly lines were obtained from Bloomington *Drosophila* Stock Center: *trpml^1^* (28992), UAS-trpm^RNAi^ (57871), *matα4-GAL-VP16* (7062), *20XUAS-IVS-GCaMP6s-attP40* (42746), *UAS-Cas9-attP2* (54595), *yw, nos-phiC31; PBac{attP-9A}* (35569).

### Buffer reagents, drug and inhibitor treatment

Isolation buffer (IB) was made as previously described (7). We found that modified Robb’s Buffer (RB) functioned as well as activation buffer (AB; (7)) for activating eggs, so we used it for this purpose in our experiments. RB contains 55mM NaOAc, 8mM KOAc, 20mM sucrose, 0. 5mM MgCl_2_, 2mM CaCl_2_, 20mM HEPES in ddH_2_O, and is adjusted to pH 6.4 with NaOH. For tests of Trpm inhibitors, stock solutions of MgCl_2_ (Mallinckrodt) and NS8593 (Sigma-Aldrich) were prepared and added to IB and RB to the indicated final concentrations, on the day of the experiment. Published concentration of Mg^2+^ (10 mM) was used to inhibit *Drosophila* Trpm (21). For NS8593, we tried a series of concentrations starting from published results of mammalian TRPM7 (30 μM) (22) and found 100 μM is the optimal concentration for *Drosophila* Trpm inhibition.

Stage-14 oocytes, dissected from virgin females that had been aged on yeast for 3-5 days, were incubated in inhibitor-treated IB or control IB for 30 min before experiments. Inhibitor-treated oocytes were then switched to incubation in RB also containing the inhibitor at the same concentration, and oocytes incubated in control IB were activated in control RB lacking the inhibitor.

### Imaging

For calcium wave visualization, oocytes were imaged using a Zeiss Elyra Super Resolution Microscope with a 5X or 10X lens with Zen software. The detection wavelength was set to 493-556 nm for GCaMP signal.

For immunostaining, fixed and stained embryos were imaged using a Zeiss LSM880 Confocal Multiphoton Microscope with a 10X lens and the Zen software. The detection wavelength was set to 495-634 nm for FITC signal, and 410-495 nm for DAPI. Z-stack images were taken from the shallowest to the deepest visible planes with a pinhole of 100. Maximum intensity projection of captured images was performed using the Zen software.

To examine ovary morphology, ovaries were dissected from 3 to 5-day old virgin females aged on yeast. Their images were captured using an Echo Revolve Microscope, with a 4X lens and brightfield imaging settings. Ovary images were processed with the Echo Revolve App.

All acquired images were processed with ImageJ software (41) as needed.

### *In vitro* egg activation assay

Oocytes were dissected from the indicated female flies and induced to activate *in vitro* following methods as described previously (7). For imaging, oocytes were placed in a drop of IB in a glass-bottomed Petri dish. After imaging parameters were configured, IB was replaced by RB to induce egg activation (9). Time-lapse images were taken at 1s or 10s intervals, for 10-20 min after the addition of RB. The distance traveled by the wavefront was measured using ImageJ software, and the elapsed time were recorded. Propagation rate of the calcium wave was calculated as distance traveled by the wavefront divided by time.

### Egg-laying and egg hatchability assay

Virgin females of the indicated genotypes were aged on yeasted food vials for 3 to 5 days, before mating with Oregon-R-P2 (ORP2) wildtype males. Matings were observed and the males were removed after a single mating had completed. Females were allowed to lay eggs in the mating vial for 24 hours and were then transferred to a new vial. Females were transferred three times before they were discarded. The number of eggs and pupae were counted. Egg hatchability was calculated by the number of pupae divided by the number of eggs. To confirm there was no post-embryonic developmental arrest in eggs laid by mutants that could have affected our hatchability score, we counted the number of unhatched eggs for 3 days after egg laying and thus determined the number of hatched eggs. This number equaled the number of pupae, indicating that our method of calculating egg hatchability was reliable.

### RT-PCR

RNA was extracted from 4-5 pairs of ovaries from virgin females aged in yeasted vials for 3-5 days using Trizol/chloroform. 750 ng RNA from each sample underwent DNase (Promega) treatment and cDNA synthesis using SuperScript™ II Reverse Transcriptase (Invitrogen) following the manufacturer’s instructions. A 30 cycle PCR amplification was performed with GoTaq polymerase (Promega) with the following conditions: 95°C 2 minutes, 95°C 40 seconds, 54°C 40 seconds, 72°C 40 seconds, 72°C 10 minutes. PCR products were run on 1% agarose gels, and the DNA was stained with 1 μg/mL ethidium bromide. Captured gel images were processed with ImageJ software.

Expression level of ribosome protein gene *rpl32* was used as a loading control (42). The following primers are used for RT-PCR: *trpm-F:* TCACTGTGCTGGTGAAGATG; *trpm-R:* CCAGAGGTCCCAGGTATTTATTC (Amplicon size with cDNA as template is 324 bp. The reverse primer spans an exon-exon junction and will amplify a 513 bp band with genomic DNA as template). *rpl32-F:* CACCAGTCGGATCGATATGC; *rpl32-R:* CGATCCGTAACCGATGTTG (Amplicon size with cDNA as template is 120 bp. The primers flank an exon-exon junction and will amplify a 182 bp band with genomic DNA as template).

### Immunostaining

Embryos and eggs from the indicated mating were collected from grape juice/agar plates. Embryos were dechorionated in 50% commercial bleach, fixed in methanol/heptane (43), and stored at 4°C until use. Fixed embryos were washed with PBST 3 times for 5 minutes each and blocked with PBST containing 5% vol of normal goat serum (PBST-NGS). Embryos were then incubated overnight at 4°C in PBST-NGS containing mouse monoclonal anti-α-Tubulin-FITC antibody (Sigma-Aldrich) at a dilution of 1:200, and RNaseA (Roche Applied Science) at a concentration of 5 μg/mL. Embryos were then washed with PBST 3 times for 5 minutes each. DNA was stained with 1 μg/mL DAPI (Molecular Probes) in PBST for 5 minutes, and embryos were mounted on glass microscope slides in anti-fade mounting buffer.

### Bleach resistance assay

Mature oocytes were dissected from indicated females in IB. Eggs laid by indicated females after mating to indicated males were collected from grape juice/agar plates. Both oocytes and embryos were incubated in 50% commercial bleach for 2 min (8). Numbers of oocytes and embryos before and after incubation were counted to calculate the percentage that survived bleach treatment.

### Statistics

Pearson’s chi square test was used to compare the frequency of calcium wave initiation and ovary morphology defects. Student’s *t* test was used to compare the propagation speed of calcium wave and egg hatchability.

## Acknowledgments

Imaging data were acquired on microscopes in the Cornell University Biotechnology Resource Center, with NSF1428922 funding for the shared Zeiss Elyra Super Resolution Microscope, and NYSTEM (CO29155) and NIH (S10OD018516) funding for the shared Zeiss LSM880 Confocal Multiphoton Microscope.

We thank Dr. C. Han for kindly providing U63-tRNA-gRNA1-tRNA-gRNA2 and CR7T-gRNA1-U63-gRNA2 plasmids and for advice on genome editing in *Drosophila* and Dr. R. Fissore for technical advice on the use of NS8593. We thank NIH grant R21-HD088744 for supporting this work. We thank Drs. C. Han, J. Liu and C. Williams, and the Wolfner lab for helpful comments on the manuscript and/or during this study.

## Supplemental Material and Methods

### Mutant verification

The deletion in *pain^TMΔ^* allele was confirmed by PCR and sequencing using the following primers: *pain-C1* F: GTATCACCACAAACGGAGAGAG; *pain-C1* R:

GGTGCCACTTGAGGAATAGAA; pain-C2F: ACCAACGTCCTTAGTCGGTA; pain-C2R: GTCCTCATTTCGAAGGGATCG (**Fig.S1A-C**).

*trpml^1^* carries a 1097bp deletion which removes nucleotides −456 to +641 relative to the *trpml* translation start site (18). The deletion was confirmed with the following primers: *trpml-F:* CTGACCACGATGTTTATCGC; *trpml-R:* AACGGCGGAATTTGGAATAC (**Fig. S1D**).

A 32 cycle PCR amplification was performed with GoTaq polymerase (Promega) with the following conditions: 95°C 2 minutes, 95°C 40 seconds, 54°C 40 seconds, 72°C 1 minute 30 seconds, 72°C 10 minutes. PCR products were run on 1% agarose gels, and the DNA was stained with 1 μg/mL ethidium bromide. Captured gel images were processed with ImageJ software.

### Single oocyte RT-PCR

RNA was extracted from single mature oocytes from virgin *trpm* germline knockout females *(nos-Cas9-attP2/gRNA-trpm^1^*) aged in yeasted vials for 3-5 days using Trizol/chloroform and underwent DNase (Promega) treatment and cDNA synthesis using SuperScript™ II Reverse Transcriptase (Invitrogen) following the manufacturer’s instructions. Three rounds of nested PCR amplification (30 cycles/round, product of the last round was used as the template for the next round) was performed with iProof Hi-Fidelity polymerase (Bio-Rad) with the following conditions: 98°C 30 seconds, 98°C 10 seconds, 56°C 30 seconds, 72°C 15 seconds, 72°C 10 minutes. The first round of PCR was performed using outer primers: trpm-outerF: GGAGTTCGAGAGCAAAGGTAAG; trpm-outerR: GAACGATCCGAGTCATCATTGT. The second and the third round of PCR were performed using inner primers: trpm-innerF: GAGCTACACTCCTGGTCAAATC; trpm-innerR: GT CAAGCT GCCT GAT GT AGAA. The final round of PCR product was run on 1 % agarose gels, and the DNA was stained with 1 μg/mL ethidium bromide. Captured gel images were processed with ImageJ software. The product bands were cut out and purified with gel extraction kit (Invitrogen) for sequencing (**Fig. S2**).

## Supplemental Figures, Movies and Tables

**Fig.SI.**
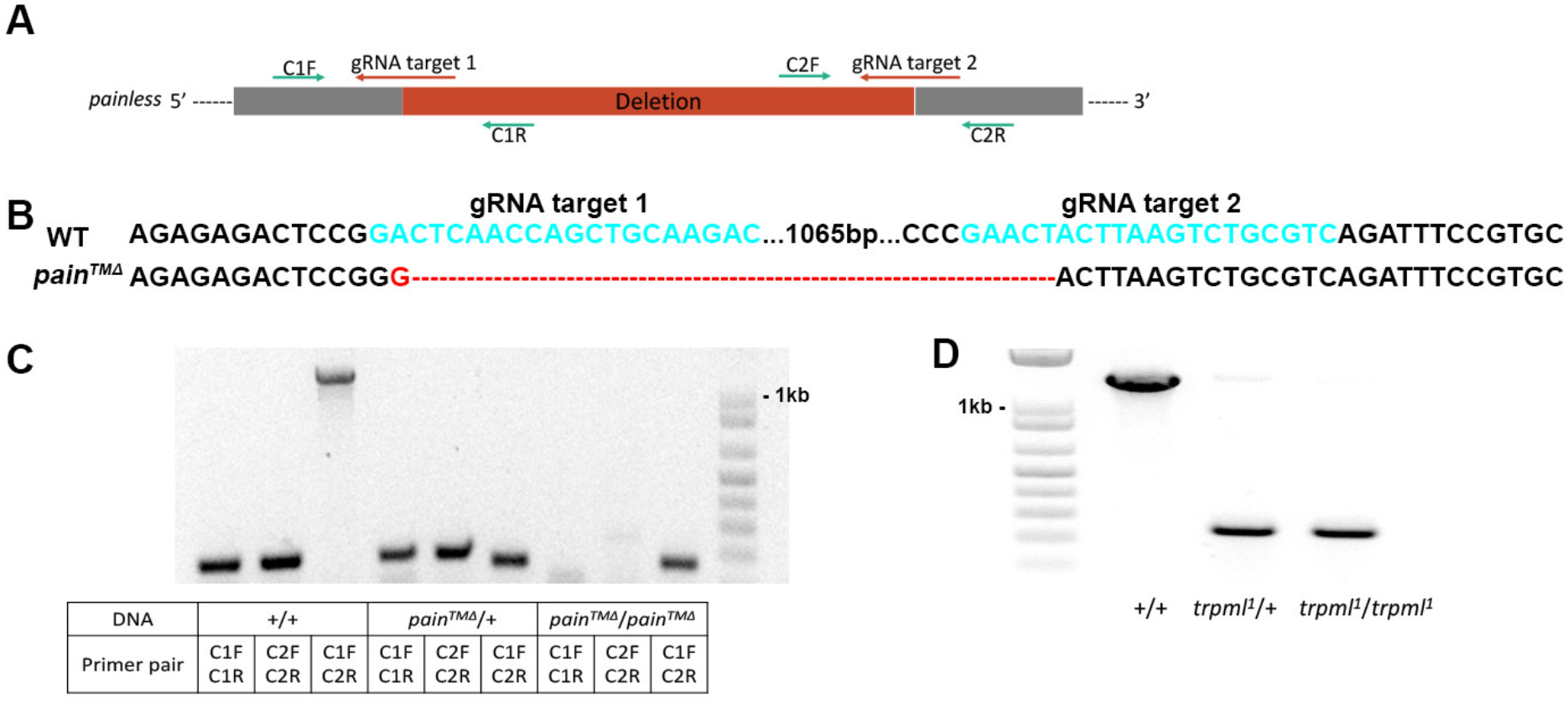
Verification of *pain* and *trpml* null mutant. **(A)** Scheme of *pain™^Δ^* deletion. Locations of gRNA targets and PCR primers relative to the deletion are shown; **(B)** Sequencing results of *pain^TMΔ^* allele. Bases labeled cyan are gRNA targets in the wildtype allele. Red labels mismatching bases and deletion resulted from CRISPR/Cas9 mediated editing; **(C)** PCR verification of *pain^TMΔ^* allele. Expected amplicon sizes: C1F+C1R (207bp in WT allele, none in mutant allele), C2F+C2R (203bp in WT allele, none in mutant allele), C1F+C2R (1268bp in wildtype allele, 177bp in mutant allele); **(D)** PCR verification of *tprml^1^* allele. Expected amplicon sizes: 1347bp in wildtype allele, 250bp in mutant allele.

**Fig.S2.**
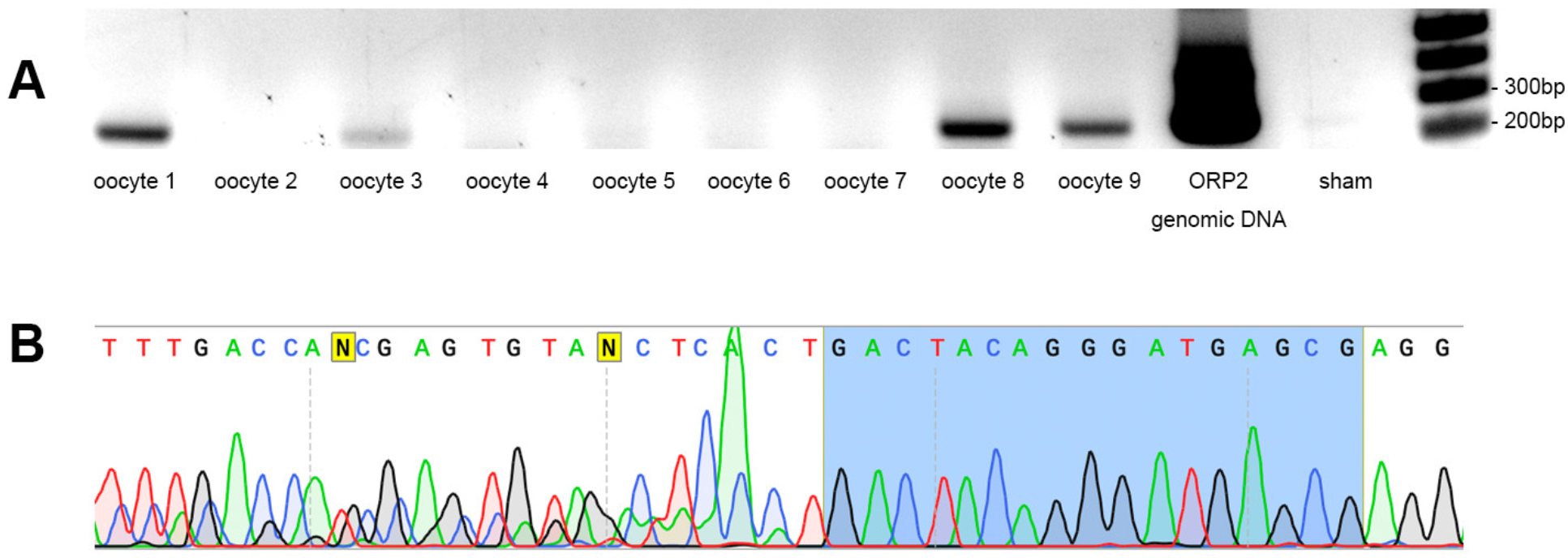
Verification of CRISPR/Cas9 generated indels in oocytes from *trpm* germline knockout females. **(A)** Representative gel of final products from single oocyte RT-PCR followed by nested PCR using oocytes from *trpm* germline knockout females *(nos-Cas9-attP2/ gRNA-trpm^1^)*. Amplicon size of *trpm* inner primers is 256bp. Oocyte 1, 3, 8 and 9 successfully generated PCR products. Oocyte 2, 4, 5, 6 and 7 failed to generate PCR products likely due to low copy number of RNA in a single egg. Genomic DNA of ORP2 wildtype flies was used as positive control. A sham sample undergoing the same RNA extraction, RT-PCR and nested PCR process was used as a negative control; **(B)** Representative sequencing results of the PCR product in **(A)**. Highlighted bases are one of the gRNA targets in gRNA set 3. Mixed sequencing peaks showed up after the gRNA target in 6 out of 8 oocyte samples sequenced, suggesting successful introduction of indels near the gRNA target by CRISPR/Cas9. We were unable to determine whether these indels caused frameshift, because it was difficult to untangle the mixed readings: depending on the timing of the editing, multiple species of *trpm* RNA can be present in an oocyte.

**Fig.S3.**
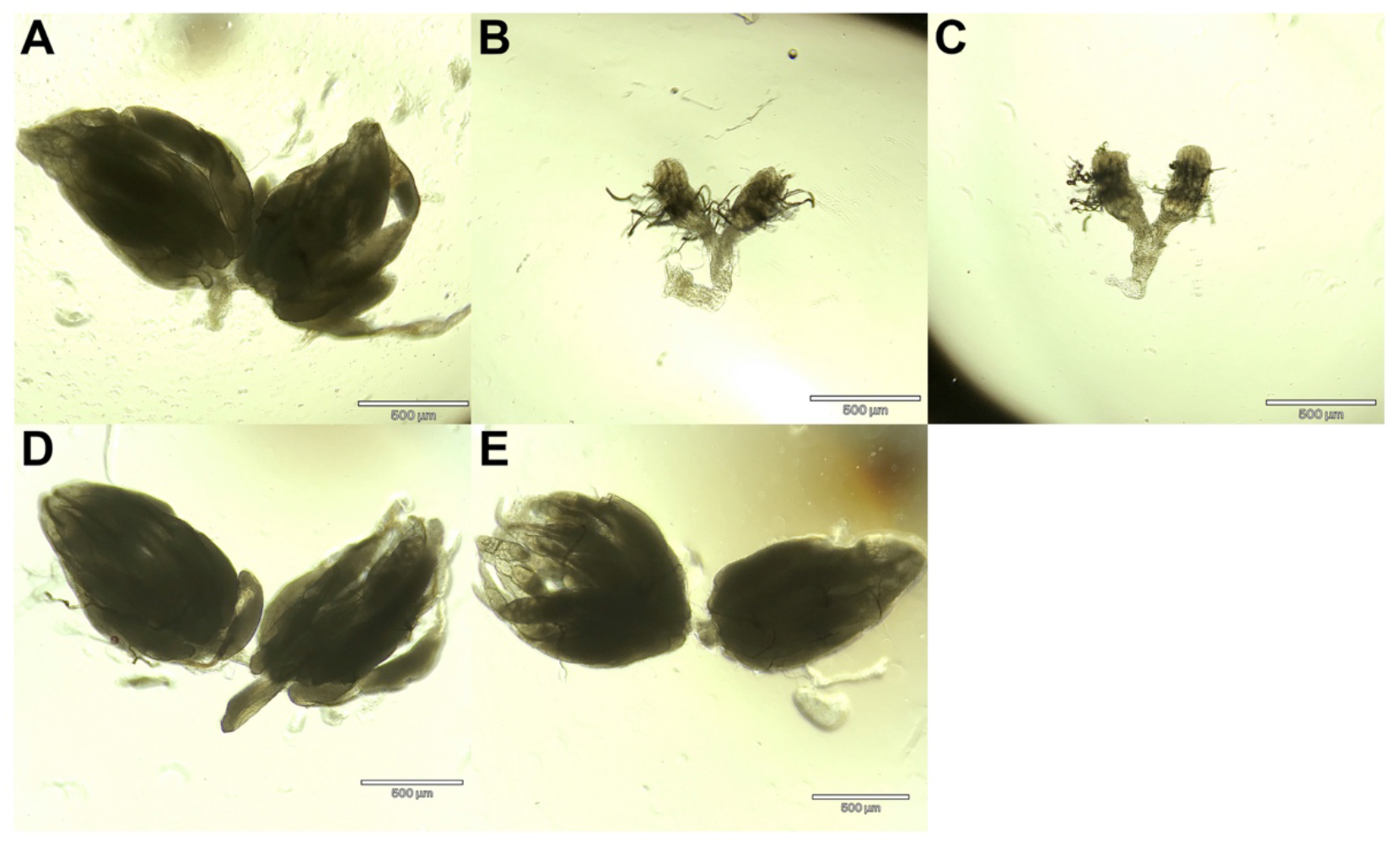
Ovary morphology of germline specific *trpm* knockout females. **(A)** *nos-Cas9* control females display normal ovary morphology (representative picture of n=25); **(B)** 34% of the *nos-Cas9; gRNA-trpm^1^* females display ovary morphology defects (representative picture of n=13/38, p=0.003); **(C)** 96% of the *nos-Cas9; gRNA-trpm^2^* females display ovary morphology defects (representative picture of n=22/23, p=2.18×10^−8^); **(D)** *matα4-GAL-VP16 > UAS-Cas9* control females display normal ovary morphology (representative picture of n=11); **(E)** *matα4-GAL-VP16 > UAS-Cas9; gRNA-trpm^1^* females display normal ovary morphology (representative picture of n=18). All scale bars=500 μm.

**Fig.S4.**
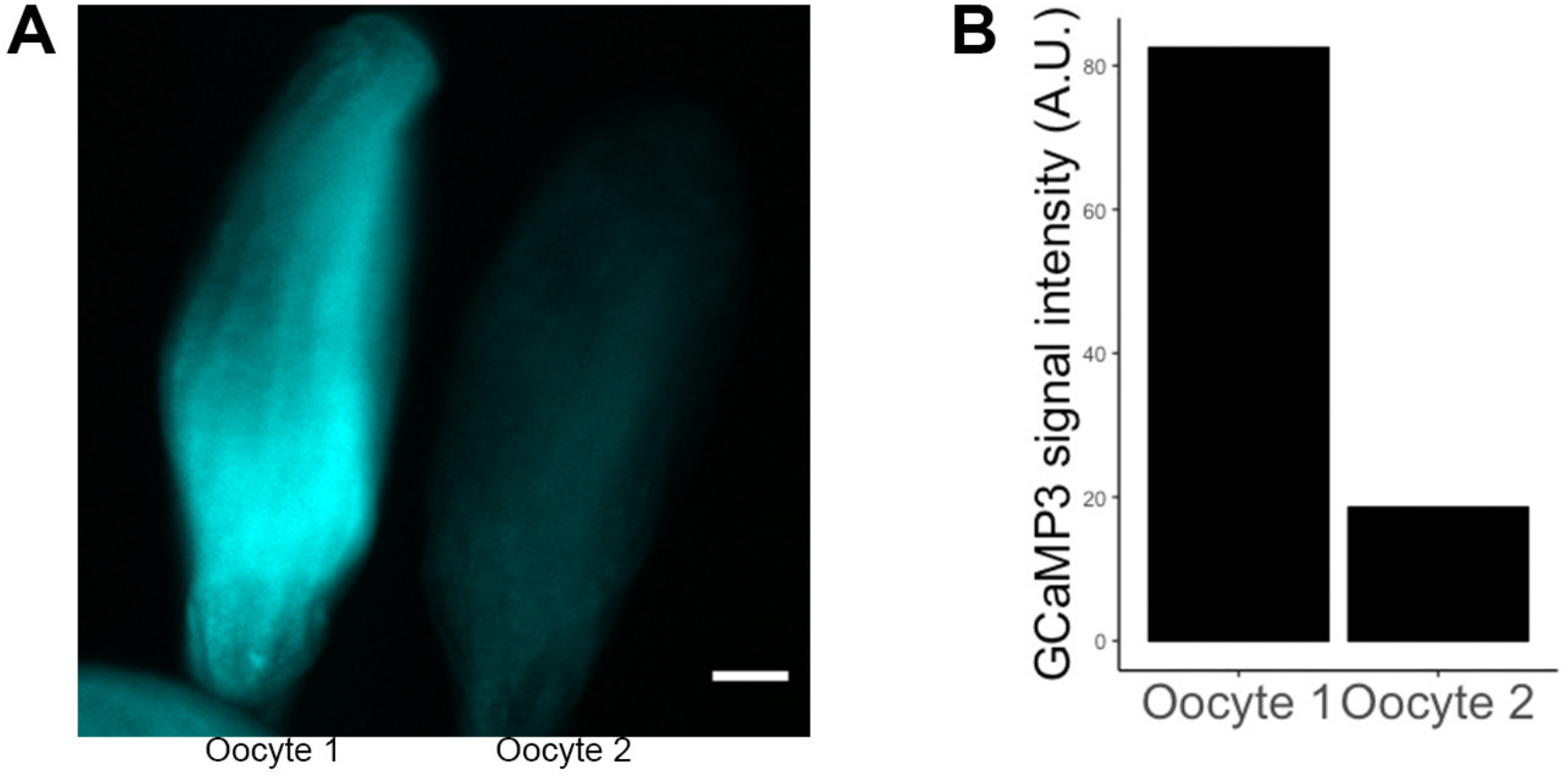
Variation in basal calcium levels in mature oocytes without a calcium wave. **(A)** Two mature oocytes dissected from the same 3 to 5-day old *matα4-GAL-VP16; UAS-GCaMP3* virgin female, incubated in IB. Difference in GCaMP3 signal suggests variation in basal calcium levels in mature oocytes. Scale bar = 50 μm; **(B)** Quantification of GCaMP3 signal intensity in **(A)**.

**Table.S1.** Efficiency test of gRNA sets targeting *trpm* using “Cas9-LEThAL” method. *act-Cas9, lig4*- mutant females were crossed to males carrying U6:3-tRNA-gRNA1-tRNA-gRNA2 plasmid with different sets of gRNAs. Number of female and male progenies from the crossing were counted. Higher gRNA efficiency results in higher male lethality.

| Efficiency test of gRNA sets targeting *trpm* using “Cas9-LEThAL” method |  |  |  |
| --- | --- | --- | --- |
| gRNA-*trpm* | Number of female progenies | Number of male progenies | Male/female ratio |
| Set 1 | 65 | 6 | 0.092 |
| Set 2 | 72 | 4 | 0.056 |
| Set 3 | 68 | 0 | 0 |

Scale bars for all movies are 50 μm.

**Movie S1**

Heterozygous *pain^TMΔ^* control oocytes calcium wave, detected with nos-GCaMP6m.

24FPS

300 frames

1 sec per frame

5 min in total

**Movie S2**

Homozygous *pain^TMΔ^* oocytes calcium wave, detected with nos-GCaMP6m.

24FPS

300 frames

1 sec per frame

5 min in total

**Movie S3**

Heterozygous *trpml^1^* control oocytes calcium wave, detected with nos-GCaMP6m.

24FPS

300 frames

1 sec per frame

5 min in total

**Movie S4**

Homozygous *trpml^1^* oocytes calcium wave, detected with nos-GCaMP6m.

24FPS

300 frames

1 sec per frame

5 min in total

**Movie S5**

Calcium wave is blocked when oocytes were treated with 10mM Mg^2^+, detected with *matα4-GAL-VP16 > UASp-GCaMP3*.

10FPS

135 frames

10 sec per frame

22.5 min in total

**Movie S6**

Calcium wave is blocked when oocytes were treated with 100μM NS8593, detected with *matα4-GAL-VP16 > UASp-GCaMP3*.

24FPS

450 frames

1 sec per frame

7.5 min in total

**Movie S7**

Calcium wave is blocked in *trpm* germline specific RNAi knockdown oocytes, detected with *matα4-GAL-VP16 > UASp-GCaMP3*.

10FPS

120 frames

10 sec per frame

20 min in total

**Movie S8**

Calcium wave is blocked in *trpm* germline specific CRISPR/Cas9 knockout oocytes with *nos-Cas9* and *gRNA-trpm^1^*, detected with *matα4-GAL-VP16 > UASp-GCaMP3*.

24FPS

360 frames

1 sec per frame

6 min in total

**Movie S9**

Calcium wave is blocked in *trpm* germline specific CRISPR/Cas9 knockout oocytes with *matα4-GAL-VP16 > UAS-Cas9* and *gRNA-trpm^1^*, detected *with matα4-GAL-VP16 > UASp-GCaMP3*.

24FPS

600 frames

1 sec per frame

10 min in total

